# mTOR disrupts cerebrovascular RNA regulatory networks in Alzheimer’s disease

**DOI:** 10.64898/2026.09.15.750223

**Authors:** Haneen Makhlouf, Andy Banh, Carlos Pomilio, Beatriz Ferran, Stacy Hussong, Flavia Saravia, Arlan Richardson, Veronica Galvan

## Abstract

Cerebrovascular dysfunction is increasingly recognized as a central contributor to the initiation and progression of Alzheimer’s disease (AD). The mammalian/mechanistic target of rapamycin (mTOR) pathway has emerged as a key driver of AD-related vascular dysfunction. Consistent with this role, attenuation of mTOR signaling restores multiple aspects of cerebrovascular function in AD models, including vascular density, cerebral blood flow, vascular reactivity, and neurovascular coupling. However, the molecular mechanisms linking mTOR dysregulation to cerebrovascular dysfunction in AD remain poorly understood we assessed cerebrovascular levels of RNA-regulatory proteins in hAPP(J20) mice treated with rapamycin from 6 to 12 months of age. Rapamycin partially restored disease-suppressed heterogeneous nuclear ribonucleoprotein (hnRNP) family members, including hnRNP D1 (AUF1) and hnRNP E1 (PCBP1). Rapamycin-mediated restoration of hnRNP extended to the cerebrovasculature of PS19 (P301S) tauopathy mice, indicating that mTOR-dependent hnRNP suppression is shared across amyloid- and tau-driven models of AD. Consistent with these findings, hnRNP E1 and hnRNP L each showed a nominally significant inverse relationship with Braak stage in AD patient brains, with a similar but non-significant trend for hnRNP R and hnRNP M; The coordinated pattern of decline suggests network-level disruption of RNA-binding protein homeostasis in AD. Together, our studies uncover a previously unrecognized link between mTOR signaling and the regulation of RNA-regulatory proteins and identify a previously unrecognized, reversible mTOR-dependent RNA regulatory program in the cerebrovasculature that may contribute to neurovascular dysfunction in AD and reveal new targets for therapeutic intervention.

## Introduction

Alzheimer’s disease (AD), the most common cause of dementia worldwide, represents a growing global health challenge. Despite extensive research, broadly effective disease-modifying therapies remain unavailable. Increasing evidence points to cerebrovascular dysfunction as an early and central contributor to AD initiation and progression [1],[2],[3]. Various vascular abnormalities, including reduced blood flow, disruption of the blood-brain barrier (BBB), endothelial dysfunction, and impaired neurovascular coupling, are observed in patients and animal models of AD and often precede abnormal neuronal loss and cognitive decline.[4],[5],[6],[7],[8]. Indeed, vascular contributions to AD are recognized as a major early driver of disease and may constitute the most clinically addressable aspect of AD. [2] The hAPP(J20) mouse model, a well-established and widely used model of AD amyloidopathy that recapitulates vascular defects of the disease, including but not limited to decreased cerebral perfusion and blood-brain barrier (BBB) disruption [9]. The mammalian target of rapamycin (mTOR) pathway is a vital regulator of cell growth, metabolism, and aging. We previously demonstrated that mTOR is a central mediator of vascular dysfunction in AD [10],[11],[12],[13],[14] and that mTOR signaling is hyperactivated in AD [10],[15]. Specifically, we showed that pharmacological suppression of mTORC1 with rapamycin restores cerebral blood flow, maintains BBB integrity, promotes amyloid clearance, and improves cognitive function in this model and in normative aging [16],[17],[18],[19],[20],[21],[22]. mTOR may thus be a key link connecting aging biology to cerebrovascular decline in AD [23]. However, the molecular effectors downstream of mTOR that cause vascular dysfunction in AD remain poorly understood.

RNA-binding proteins and post-transcriptional regulatory mechanisms are increasingly recognized as contributors to neurodegenerative diseases. Large-scale proteomic and transcriptomic analyses of AD brain tissue have identified RNA-binding proteins and RNA-splicing modules as among the most disease-associated molecular signatures, strongly linked to neuropathology and cognitive decline [24],[25],[26],[27],[28]. Changes in RNA processing, ribonucleoprotein dynamics, and translational control have been observed in AD and related disorders [29],[30]. Yet it remains unknown whether RNA regulatory networks within the cerebrovasculature undergo remodeling in AD, and whether these changes are regulated by mTOR signaling. Heterogeneous nuclear ribonucleoproteins (hnRNPs) regulate multiple aspects of RNA metabolism, including pre-mRNA splicing, mRNA stability, transport, and translation [29],[30],[31],[32],[33]. Within this family, hnRNP D (AUF1), hnRNP E1 (PCBP1), and hnRNP AB modulate the stability and translation of transcripts involved in inflammatory and stress-responsive pathways [34],[35],[36],[37],[38]. HNRNP D1 and HNRNP E1 have been implicated in aging and neurodegenerative processes, linking RNA regulatory control to cellular senescence and proteostasis decline [37],[38],[39]. Given that vascular aging and mTOR signaling are central contributors to AD pathogenesis, suppression of these RNA-binding proteins may represent a downstream mechanism linking mTOR dysregulation to cerebrovascular dysfunction. Consistent with this, mTOR attenuation restored several RNA-processing proteins within the cerebrovascular proteome, identifying an mTOR-sensitive RNA regulatory axis in AD vasculature. Here, we identify a previously unrecognized mTOR-dependent RNA regulatory mechanism in the cerebrovasculature that contributes to neurovascular dysfunction in AD and can be reversed by rapamycin, and potentially other interventions reducing signaling through mTOR.

## Methods

### Animals

The derivation and characterization of hAPP(J20) transgenic mice, a widely used model of amyloidopathy of Alzheimer’s disease (AD), have been described previously [40]. Briefly, hAPP(J20) mice express a minigene encoding the human amyloid precursor protein (hAPP) with all its splicing sites carrying the familial AD– associated Swedish (K670N/M671L) and Indiana (V717F) mutations, under the control of a platelet-derived growth factor-β (PDGF-β) neuron-specific promoter active during embryonic stages. Heterozygous hAPP(J20) mice were maintained by heterozygous breeding with C57BL/6J mice, and non-transgenic wild-type (WT) littermates were used as controls. For proteomics experiments, mice were randomly assigned to receive either chow containing encapsulated rapamycin (14 ppm) or control chow containing the encapsulation material Eudragit alone, beginning at 6 months of age and continuing until 12 months of age. The cohort used for Wes’s capillary immunoassay validation experiments had received the same doses of encapsulated rapamycin or Eudragit from 10 to 12 months of age.

At the end of treatment, mice were euthanized under isoflurane anesthesia followed by cervical dislocation, and brain tissues were rapidly collected and flash-frozen for downstream analyses. Brain tissues from PS19 (P301S) transgenic mice [41], and age-matched controls were collected and processed for capillary immunoassay using identical protocols. All animal procedures involving J20 mice were approved by the Institutional Animal Care and Use Committee (IACUC) at the University of Texas Health Science Center at San Antonio (UT Health San Antonio) and were conducted in accordance with the National Institutes of Health Guide for the Care and Use of Laboratory Animals. Procedures involving PS19 mice were approved by the Institutional Animal Care and Use Committee (IACUC) at the University of Oklahoma Health Sciences Center and conducted in accordance with these guidelines.

### Cerebral blood flow measurements and analyses

Evoked cerebral blood flow (CBF) responses were measured in PS19 transgenic (Tg) and non-transgenic (NTg) mice as described in [20]. Animals were divided into three groups: NTg treated with control (Eudragit) chow (WT; n = 3), Tg treated with control (Eudragit) chow (PS19+Veh; n = 3), and Tg treated with rapamycin chow (PS19+Rapa; n = 4). Changes in CBF were recorded during sensory stimulation and expressed as a fold change from baseline. Time-course responses were averaged within each group and shown as mean ± SEM. The stimulation period is indicated on the time axis. To quantify the hemodynamic response, the area under the curve (AUC) during the stimulation window was calculated for each animal. Group differences in AUC were analyzed using one-way ANOVA followed by Tukey’s HSD post hoc test for multiple comparisons.

### Postmortem human brain tissue

Samples were obtained from the Michigan Brain Bank. The cohort comprised 22 individuals (11 male, 11 females; age range 59–90 years; PMI range 3–24 hours), including 7 neurologically normal controls (Braak stage N/A) and 15 individuals with neuropathological AD diagnoses spanning Braak stages IV–VI. Braak staging was performed by the Michigan Brain Bank according to standard neuropathological criteria. Somatosensory cortex was used for all protein quantification; per-protein n values ranged from 19 to 21 depending on antibody signal detection. Frozen brain samples were homogenized in liquid nitrogen, lysed by sonication in 1× Cell Lysis Buffer (Cell Signaling Technology, #9803) supplemented with protease inhibitor cocktail (complete Mini, Roche), and protein concentration was determined by Bradford assay (Bio-Rad).

### Isolation of brain microvasculature

Mice were euthanized under isoflurane anesthesia followed by cervical dislocation. Frozen brains were minced into approximately 1 mm^3^ pieces and homogenized in MCDB131 medium (Gibco, Waltham, MA, USA) supplemented with 2% Cosmic calf serum (CCS; HyClone, Logan, UT, USA) using a loose-fitting 7 mL Dounce homogenizer. Homogenized samples were mixed with an equal volume of MCDB131 medium containing 2% CCS and dextran (70,000 molecular weight; Sigma-Aldrich, St. Louis, MO, USA) to a final dextran concentration of 17% (w/v) and centrifuged at 12,348 × g for 15 minutes at 4 °C. The resulting pellets containing enriched brain microvascular fractions were lysed by sonication in 1× Cell Lysis Buffer (Cell Signaling Technology, #9803, Danvers, MA, USA) supplemented with protease inhibitor cocktail (complete Mini, Roche, Indianapolis, IN, USA).

### Protein quantification

Protein expression was quantified by automated capillary western blot using the Jess Simple Western system (Bio-Techne/Protein Simple, San Jose, CA) according to the manufacturer’s protocol. Chemiluminescent signals were measured as area under the curve. Target protein signals were normalized to total protein using the Protein Simple Total Protein detection module or to β-Actin (UBP Bio, Y1050) or Vinculin as a loading control, depending on the experiment. The following primary antibodies were used: β-Actin (UBP Bio, Y1050, Y1051) HNRNP E1 (Protein tech, 14523-1-AP), hnRNP R (Novus Biologicals, NBP3-02996), hnRNP L (Novus Biologicals, NBP3-03641), hnRNP M (Novus Biologicals, NB200-315SS), hnRNP K (Novus Biologicals, NBP2-24532SS), and hnRNP A/B (Santa Cruz Biotechnology, SC-37641).

### Statistical analysis and software

All statistical analyses were performed in GraphPad Prism (v10.0.0; GraphPad Software) unless otherwise specified. Graphical visualizations were done in R (v4.5.2). For immunoblot analyses, two-sided unpaired t tests were used for comparisons between two groups. Comparisons among the three groups shown in Fig. 5I were assessed by one-way ANOVA followed by Tukey’s post hoc test.

For postmortem human brain tissue, associations between protein abundance and Braak stage were assessed using two-sided Pearson correlation tests. Neuropathologically normal controls, for whom Braak stage was not applicable, were assigned a Braak stage of 0 for these analyses. Benjamini–Hochberg false discovery rate correction was applied across the six hnRNP proteins tested and is reported as q-values. One sample was excluded from the hnRNP K correlation as a statistical outlier using the ROUT method (Q = 1%) in GraphPad Prism. No other outliers were excluded from the analyses.

## Results

Accumulating evidence [10],[15],[16],[17],[18],[19],[20],[21],[22],[42] points to mTOR as a key driver of AD-related vascular dysfunction since attenuation of mTOR signaling with rapamycin dramatically improves cerebrovascular function in AD [10],[15],[16],[17],[18],[19],[20],[21],[22],[42] and aging models [22]. The molecular mechanisms linking mTOR dysregulation to cerebrovascular dysfunction, however, remain largely unknown. To map the molecular pathways involved in mTOR-driven vascular dysfunction in AD, we performed unbiased quantitative proteomic profiling of cerebral microvessels purified from hAPP(J20) mice [43],[44], a well-defined, extensively validated model of AD, and their unaffected non-transgenic littermates. All groups were treated with rapamycin encapsulated in the chow or with vehicle [5],[10],[45],[46],[47],[48].

### Rapamycin restores hnRNP D1 (HNRNP D1) levels and isoform balance in hAPP(J20) cerebrovascular microvessels

Further analyses of hnRNP abundance revealed an age-dependent pattern for both HNRNP E1 and hnRNP D1 (AUF1) in purified hAPP(J20) mouse microvessels **(Fig. 1B, E)**. At 6 months, cerebrovascular HNRNP E1 levels did not yet differ significantly between WT and vehicle-treated hAPP(J20) mice, but rapamycin treatment significantly increased HNRNP E1 above both WT and vehicle levels **(Fig. 1C)**. By 12 months, HNRNP E1 was significantly decreased in vehicle-treated hAPP(J20) mice relative to WT and was partially restored by rapamycin **(Fig. 1D)**. hnRNP D1 showed a similarly age-dependent, but inverse, pattern: at 6 months, hnRNP D1 was significantly elevated in vehicle-treated hAPP(J20) microvessels relative to WT, and this elevation was not significantly altered by rapamycin treatment **(Fig. 1F);** by 12 months, hnRNP D1 was significantly decreased in vehicle-treated hAPP(J20) mice relative to WT and was partially restored by rapamycin **(Fig. 1G)**. These data indicate that HNRNP E1 and hnRNP D1 undergo dynamic, age-dependent regulation across disease progression, transitioning from an early, normal levels to a later, disease-associated deficit partially reversed by mTOR attenuation. Since HNRNP D1 influences inflammatory mRNA stability, we next investigated whether its dysregulation extended beyond the cerebrovasculature. HNRNP D1 exists as four alternatively spliced isoforms, p38, p40, p42, and p45, distinguished by molecular weights ranging from 38 to 45 kDa [49]. Distribution of HNRNP D1 isoforms varied between tissue types: multiple isoforms were present in whole-brain lysates, whereas purified microvessels predominantly contained the higher-molecular-weight isoforms **(Fig. 1F, H)**. In hAPP(J20) microvessels, HNRNP D1 isoforms were significantly reduced compared to WT controls but were restored after rapamycin treatment **(Fig. 1F-G)**. Of note, HNRNP D1 isoforms and total HNRNP D1 were unaffected by amyloidopathy in whole brains of hAPP(J20) mice **(Fig. 1H-I)**. Nevertheless, mTOR attenuation by rapamycin increased HNRNP D1 levels in the whole brain **(Fig. 1H-I)**. Immunofluorescence analyses showed HNRNP D1 enrichment in CD31-positive cells in cortical microvessels **(Fig. 1A)**, suggesting that HNRNP D1 levels by capillary immunoassays emanate from brain microvascular endothelium. Levels of vascular structural proteins, such as actin and vinculin, remained consistent across genotypes and treatments and served as loading controls **(Fig. 1H)**. Together, these results suggest that HNRNP D1 depletion is specific to the cerebrovascular compartment in AD-like amyloidosis and is dependent on mTOR activity, while mTOR may act broadly to regulate HNRNP D1 levels in the brain. Overall, these findings identify HNRNP D1 as a vascular-enriched RNA-binding protein that is depleted and exhibits isoform imbalance in AD models. Restoring HNRNP D1 levels and isoform balance after mTOR inhibition supports a model in which mTOR signaling contributes to cerebrovascular dysfunction by disturbing RNA-regulatory protein networks.

**Figure 1.**
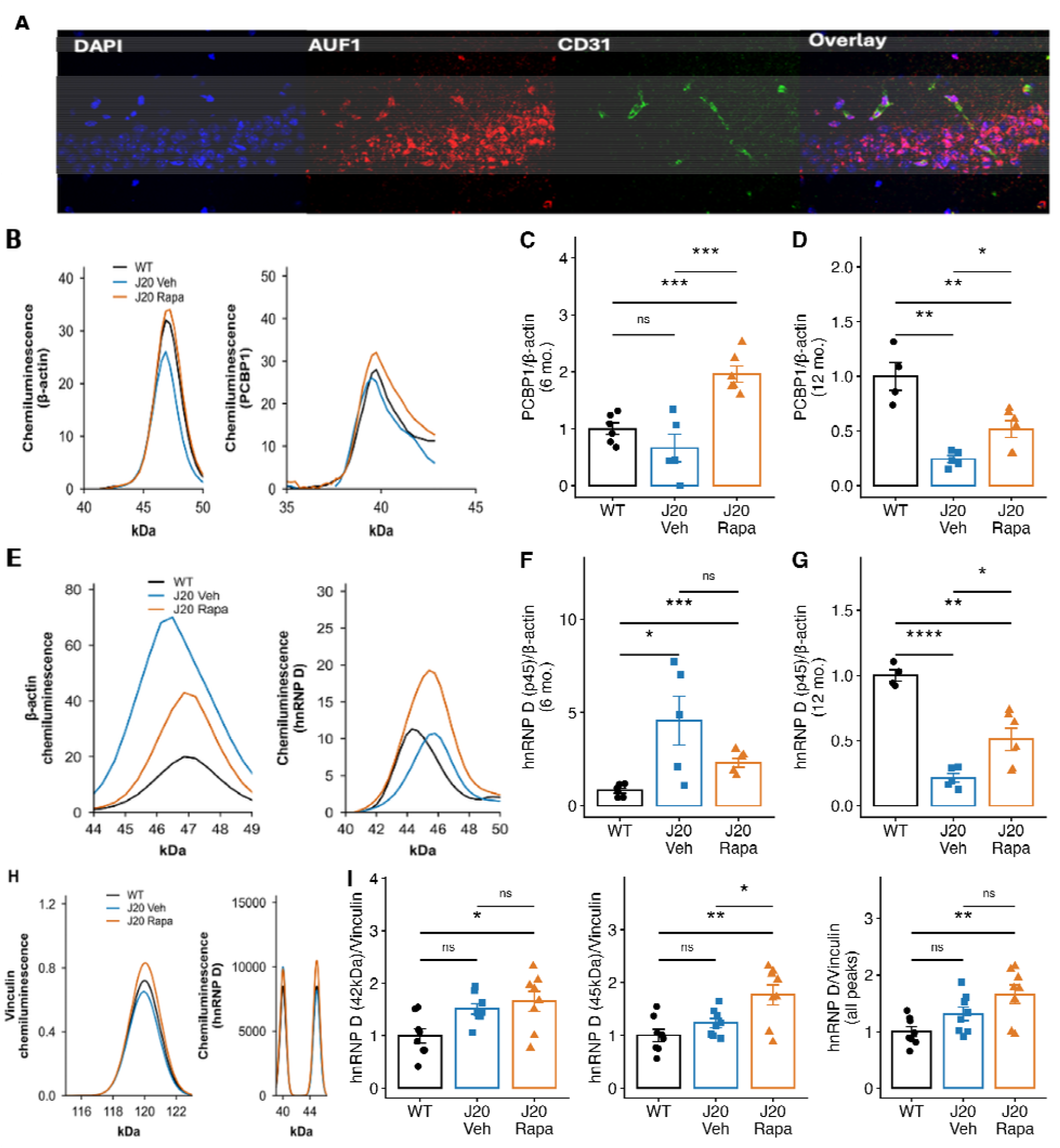
hnRNP D1 /hnRNP D and hnRNP E1 deficits in hAPP (J20) mice show rapamycin-responsive recovery selectively in the cerebrovasculature. (A) Representative immunofluorescence images of mouse brain sections stained for DAPI (nuclei), HNRNP D1 /hnRNP D, and CD31 (endothelial marker), with merged images showing HNRNP D1 localization in CD31-positive cerebrovascular structures. (B) Representative Jess capillary electrophoresis immunoassay traces for β-actin and HNRNP E1 in isolated cerebrovascular fractions. (C) Cerebrovascular HNRNP E1 signal normalized to β-actin at 6 months (WT, n□=□6; J20+Veh, n□=□5; J20+Rapa, n□=□6). WT versus J20+Veh, t (9)□=□1.38, P□=□0.201 (ns); WT versus J20+Rapa, t (10)□=□5.42, P□=□0.0003; J20+Veh versus J20+Rapa, t (9)□=□4.83, P□=□0.0009. (D) Cerebrovascular HNRNP E1 signal normalized to β-actin at 12 months (WT, n□=□4; J20+Veh, n□=□5; J20+Rapa, n□=□6). WT versus J20+Veh, t (7)□=□6.45, P□=□0.0003; WT versus J20+Rapa, t (8)□=□3.48, P□=□0.0083; J20+Veh versus J20+Rapa, t (9)□=□3.13, P□=□0.0121. (E) Representative Jess traces for β-actin and HNRNP D1 /hnRNP D in isolated cerebrovascular fractions. (F) Cerebrovascular HNRNP D1 (p45) signal normalized to β-actin at 6 months (WT, n□=□6; J20+Veh, n□=□5; J20+Rapa, n□=□6). Because Brown-Forsythe testing indicated unequal variances among groups, WT versus J20+Veh and J20+Veh versus J20+Rapa were evaluated by Welch’s t test: WT versus J20+Veh, t (4.1)□=□2.84, P□=□0.0458; J20+Veh versus J20+Rapa, t (4.3)□=□1.70, P□=□0.161 (ns). WT versus J20+Rapa was evaluated by Student’s t test: t (10) = 5.42, P = 0.0003. (G) Cerebrovascular HNRNP D1 /hnRNP D (p45) signal normalized to β-actin at 12 months (WT, n□=□4; J20+Veh, n□=□5; J20+Rapa, n□=□6). WT versus J20+Veh, t (7)□=□14.71, P□<□0.0001; WT versus J20+Rapa, t (8)□=□4.36, P□=□0.0024; J20+Veh versus J20+Rapa, t (9)□=□2.96, P□=□0.0159. (H) Representative Jess traces for vinculin and HNRNP D1 /hnRNP D isoforms in whole-brain lysates. (I) Quantification of HNRNP D1 /hnRNP D isoform-specific signals normalized to vinculin in whole-brain lysates (n□=□8 per group), including the p42 isoform (left), p45 isoform (center), and total signal across all detected isoforms (right), evaluated by one-way ANOVA with Tukey’s multiple-comparisons test. For p42, F (2,21)□=□5.40, P□=□0.013; WT versus J20+Rapa, P□=□0.014, with all other comparisons not significant. For p45, F (2,21)□=□8.10, P□=□0.0025; WT versus J20+Rapa, P□=□0.0021; J20+Veh versus J20+Rapa, P□=□0.032; WT versus J20+Veh, not significant. For total HNRNP D1 /hnRNP D, F (2,21)□=□6.75, P□=□0.0054; WT versus J20+Rapa, P□=□0.0039, with all other comparisons not significant. Individual points represent biological replicates; bars indicate means ± SEM. ns, not significant; *P□<□0.05; **P□<□0.01; ***P□<□0.001; ****P□<□0.0001.

### Progressive loss of specific hnRNP family members with increasing Braak stage in AD patient brains

hnRNP levels are associated with AD progression; we examined the relationship between Braak stage and hnRNP protein abundance. hnRNP E1 (r = −0.43, P = 0.039) and hnRNP L (r = −0.59, P = 0.005) showed significant inverse associations with Braak stage **(Fig. 2A–B)**. while hnRNP R (r = −0.34, P = 0.111) and hnRNP M (r = −0.33, P = 0.130) showed negative but nonsignificant trends **(Fig. 2C–D)**. In contrast, hnRNP K (r = −0.10, P = 0.681) and hnRNP A/B (r = −0.08, P = 0.740) showed no association with Braak stage **(Fig. 2E–F)**. These analyses were not adjusted for age, sex, PMI, or other covariates because the cohort was not powered for multivariable analyses. Together, these findings support a selective decrease in several hnRNP proteins with advancing AD pathology, rather than a universal reduction across the hnRNP family.

**Figure 2.**
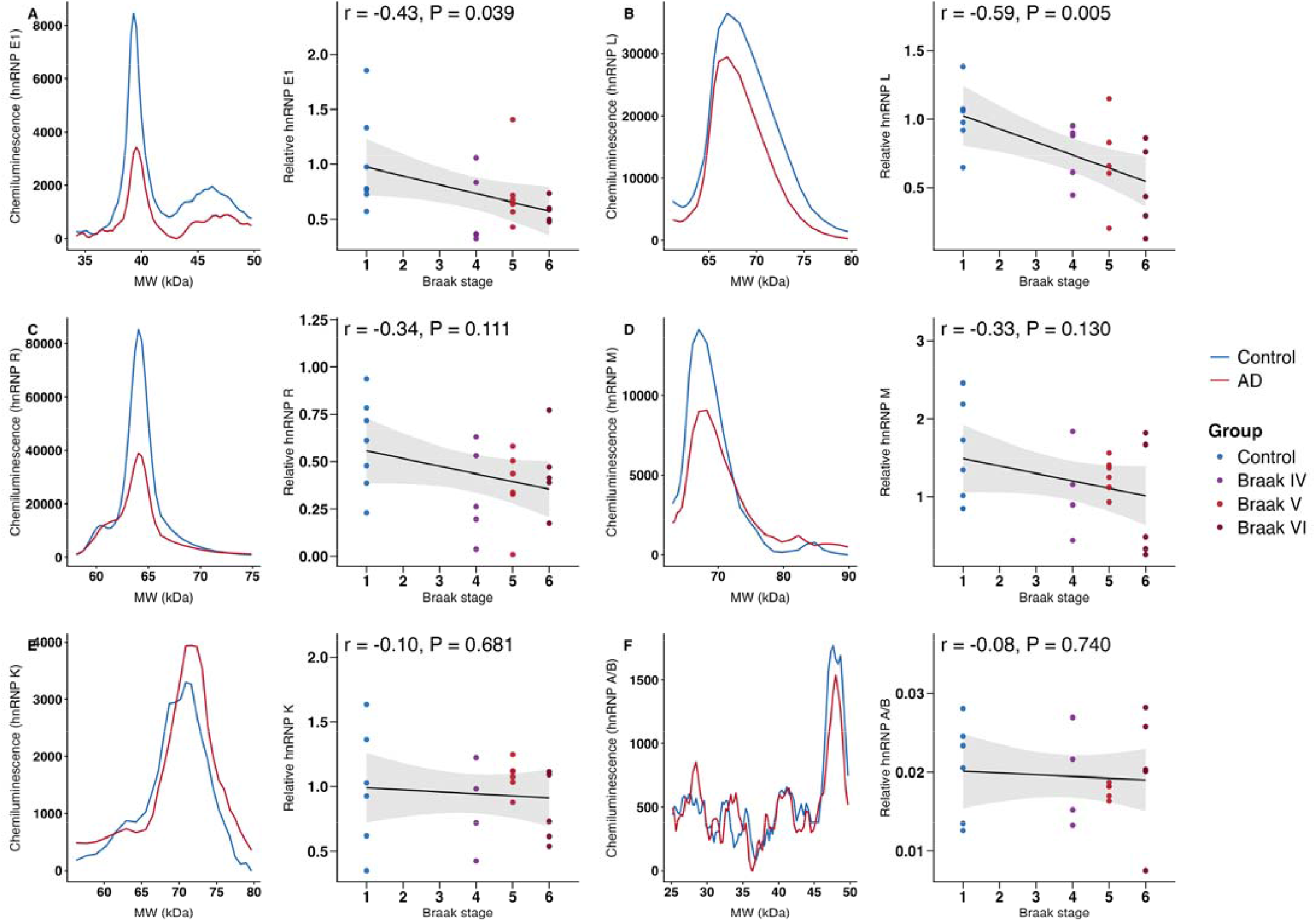
Advancing Braak neuropathology is associated with selective decline of vulnerable hnRNP RNA-binding proteins in human brain tissue. (A–F) JESS capillary immunoassay analysis of hnRNP RNA-binding proteins in human brain lysates across Braak stages. Representative electropherogram traces are shown to the left of their corresponding quantification plots. Protein abundance was normalized to vinculin and plotted by Braak stage for (A) hnRNP E1/PCBP1 (n = 23), (B) hnRNP L (n = 21), (C) hnRNP R (n = 23), (D) hnRNP M (n = 22), (E) hnRNP K (n = 21), and (F) hnRNP A/B (n = 20). Each point represents an individual human brain sample. Colors indicate Braak group. Solid lines show Pearson linear-regression fits across individual samples; shaded areas indicate 95% confidence intervals. Pearson correlation coefficients and two-tailed P values are displayed in each panel. One statistical outlier was excluded from the hnRNP K analysis using the ROUT method (Q = 1%; see Methods). hnRNP E1 and hnRNP L showed inverse associations with Braak stage.

### Rapamycin improves cerebrovascular function and restores hnRNP E1 in the PS19(P301S) model of tauopathy

We previously demonstrated that cerebrovascular functional and cognitive deficits occur in the hAPP(J20) model and that these deficits are improved by rapamycin treatment [17],[20],[21],[22]. We assessed cerebrovascular responses to somatosensory stimulation in PS19(P301S) tauopathy mice, both treated and untreated with rapamycin **(Fig. 3A)**, and in WT littermates as controls. WT mice exhibited robust increases in vascular response following stimulation, whereas PS19 (P301S) mice showed markedly reduced responses, consistent with impaired neurovascular function **(Fig. 3B)**. Rapamycin treatment significantly improved the magnitude of the vascular response in PS19(P301S) mice, partially restoring responses toward WT levels **(Fig. 3C)**. Quantification of response amplitude confirmed a trend toward decreased stimulus-evoked vascular responses in PS19(P301S) mice compared to WT controls **(Fig. 3C)**. Rapamycin treatment increased response amplitude in PS19(P301S) mice, indicating partial recovery of vascular responsiveness **(Fig. 3C)**. These findings demonstrate that mTOR attenuation enhances cerebrovascular functional responses in a model of AD tauopathy. To determine whether mTOR attenuation restores RNA regulatory proteins identified in our proteomic analysis **(Fig 1 B-H)** in the PS19(P301S) model of tauopathy, we measured hnRNP E1 levels in isolated cerebrovascular fractions from P301S animals **(Fig. 3D–E)**. Similar to our findings in hAPP(J20) mice modeling amyloidopathy, hnRNP E1 abundance was significantly decreased in the cerebrovasculature of PS19 (P301S) mice compared with WT controls (WT vs. PS19+Vehicle, P < 0.01). Rapamycin treatment significantly increased hnRNP E1 levels in PS19 mice relative to vehicle-treated PS19 mice (PS19+Vehicle vs. PS19+Rapamycin, P < 0.05), and rapamycin-treated PS19 (P301S) animals were no longer statistically distinguishable from WT controls. Together, these findings suggest that mTOR-dependent hnRNP dysregulation is not restricted to amyloid-driven pathology but extends to a tau-driven model of cerebrovascular dysfunction, consistent with a potential role for tau downstream of amyloid in mediating hnRNP depletion in the AD microvasculature.

**Figure 3.**
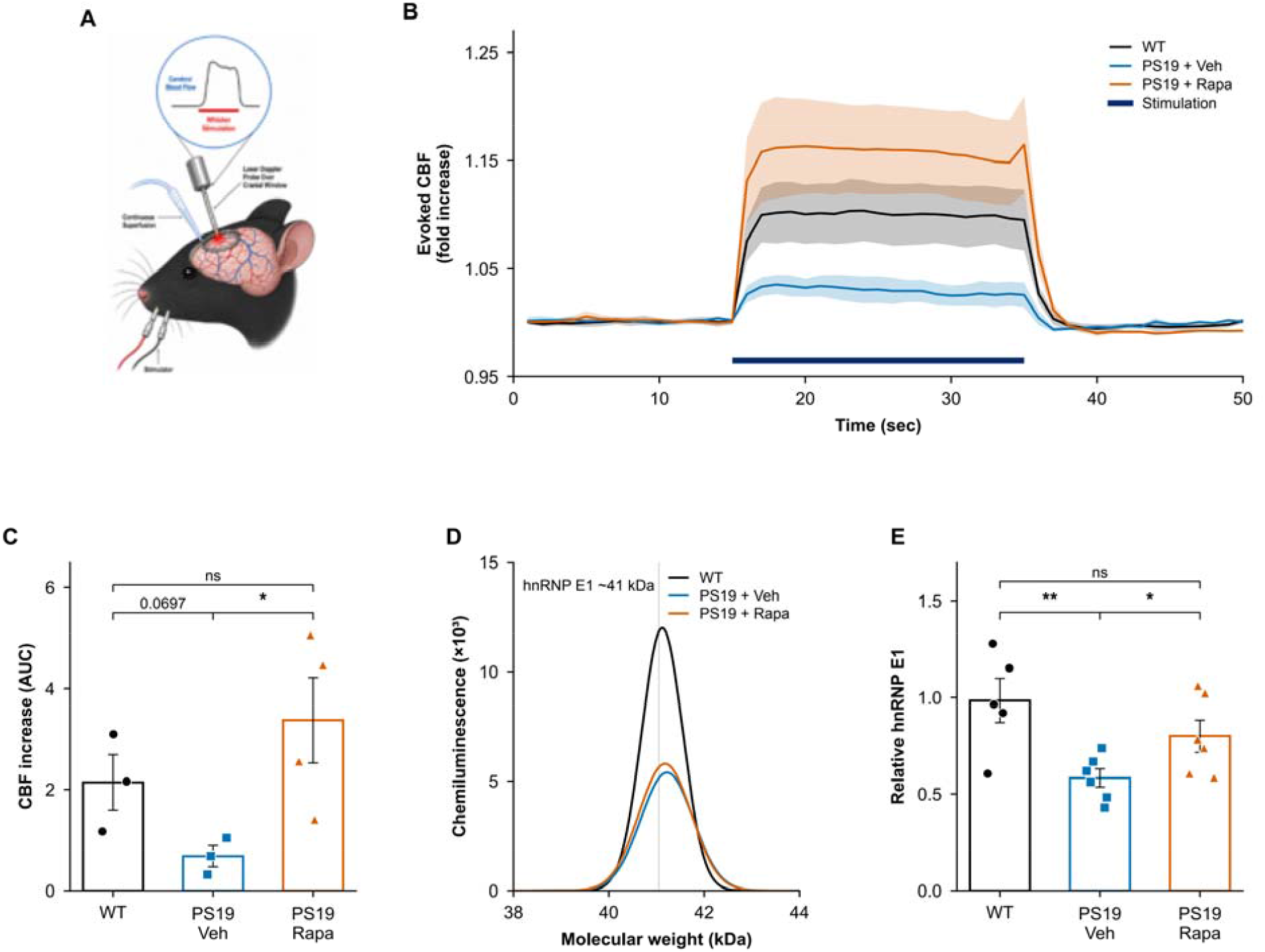
mTOR attenuation improves neurovascular coupling in the PS19 tauopathy model and partially restores hnRNP E1 levels, linking functional rescue of the vasculature to regulation of the RNA-processing pathway. (A) Schematic of neurovascular coupling assessment by laser Doppler flowmetry during whisker-pad stimulation. Cerebral blood flow (CBF) was recorded through a cranial window before, during, and after sensory stimulation. (B) Mean evoked CBF responses in WT, PS19+Veh, and PS19+Rapa mice during 20-s whisker-pad stimulation. Traces show group means ± SEM and are expressed as fold change relative to the pre-stimulation baseline. The stimulation period is indicated by a blue horizontal bar. (C) Quantification of the evoked CBF response as area under the curve (AUC) during stimulation. CBF responses were lower in PS19+Veh mice than in WT mice, although the difference did not reach statistical significance (two-tailed unpaired t test: t (4) = 2.460, P = 0.0697). Rapamycin increased the evoked CBF response in PS19 mice (PS19+Rapa versus PS19+Veh; two-tailed unpaired t test with Welch’s correction: t (3.362) = 3.092, P = 0.0461). PS19+Rapa and WT mice did not differ significantly (t (5) = 1.118, P = 0.3144). n = 3 WT, n = 3 PS19+Veh, and n = 4 PS19+Rapa mice. (D) Representative Wes capillary immunoassay electropherograms showing hnRNP E1/HNRNP E1 at approximately 41 kDa in isolated brain microvessels from (WT, PS19+Veh, and PS19+Rapa mice). (E) Quantification of hnRNP E1/HNRNP E1 abundance normalized to total protein. hnRNP E1/HNRNP E1 was reduced in PS19+Veh mice relative to WT mice (t (9) = 3.454, P = 0.0072) and increased in PS19+Rapa mice relative to PS19+Veh mice (t (10) = 2.268, P = 0.0467). PS19+Rapa and WT mice did not differ significantly (t (9) = 1.338, P = 0.2139). Two-tailed unpaired t tests were used. n = 5 WT, n = 6 PS19+Veh, and n = 6 PS19+Rapa mice. In (C) and (E), points represent individual mice, and bars show means ± SEM. *P < 0.05, **P < 0.01; ns, not significant.

## Discussion

Cerebrovascular dysfunction is increasingly identified as a key driver for the initiation and progression of AD [1],[2],[3],[4],[5],[6],[7],[8]. Early reductions in cerebral blood flow, breakdown of the blood-brain barrier (BBB), and impaired neurovascular coupling occur in patients and experimental models of AD. These cerebrovascular deficits often precede neuronal loss and cognitive decline [4],[5],[6],[50],[51]. Amyloid-β (Aβ) clearance occurs through the vascular system; compromised cerebrovascular integrity promotes Aβ accumulation and creates a feed-forward cycle in which vascular dysfunction accelerates amyloid pathology and further disrupts vascular homeostasis [3],[5],[52],[53]. Both clinical and experimental evidence support the view that vascular deficits are fundamental to disease progression rather than just secondary effects. Our [10],[15],[16],[17],[19],[20],[21],[22],[42],[46],[48],[54],[18] and others’ [55],[56],[57] prior work identified the mechanistic target of rapamycin (mTOR) signaling as a key mediator of cerebrovascular dysfunction in AD [24],[25],[27],[28],[58],[59], contributing to impaired cerebral blood flow, blood-brain barrier integrity, and neurovascular coupling that underlie AD-related vascular dysfunction [10],[16],[17],[19],[20],[21],[22],[46],[47],[18]. We previously demonstrated that mTOR signaling is aberrantly hyperactivated in AD [60], and pharmacological inhibition of mTOR by rapamycin restores cerebral blood flow, preserves BBB integrity, rescues NVC responses, and improves cognitive performance in hAPP(J20) mice [12],[16],[17],[19],[20],[21],[22],[18]. These findings agree with a growing body of evidence linking mTOR activation to vascular aging, endothelial dysfunction, impaired autophagy, and neurodegeneration [10],[20],[18].

Large-scale proteomic network analyses of AD brains have identified RNA-binding proteins and RNA-splicing enriched modules that are strongly linked to neuropathology and cognitive decline [24],[26],[27],[28]. Similarly, transcriptomic studies have shown widespread changes in RNA splicing and RNA regulatory pathways in AD brain tissue [59]. Many of these RNA regulatory modules include heterogeneous nuclear ribonucleoproteins (hnRNPs) and spliceosome components [61],[62],[63]. The selective, mTOR-dependent downregulation of snRNPs in the brain microvasculature of AD models demonstrates that disruption of RNA-regulatory networks is not limited to neurons but also involves changes in the brain microvasculature. Moreover, while hnRNPs [64],[65] and other regulatory RNA families such as lncRNAs [58],[66] have been implicated in the regulation of mTOR, the role of mTOR in the regulation of hnRNPs (except in a single report of mTOR binding to hnRNP F/H during cell proliferation [67]), and the involvement of mTOR-mediated regulation of hnRNPs in microvascular dysfunction of AD have not been previously described.

This study reveals a coordinated suppression of RNA regulatory proteins, including but not limited to hnRNPs, in the microvasculature of a model of AD amyloidosis, that were recapitulated in a model of tauopathy. We found that proteins involved in RNA processing and homeostasis were strongly suppressed in microvessels, but not in parenchyma, of AD model mice, and that this suppression could be negated by attenuation of mTOR activity. Thus, our studies suggest that previously unrecognized mTOR-dependent disruption of vascular RNA regulatory mechanisms contributes to neurovascular dysfunction in AD and can be reversed by mTOR inhibition with rapamycin.

Heterogeneous nuclear ribonucleoproteins (hnRNPs) were prominent components of the set rescued in microvasculature of rapamycin-treated AD mice. The hnRNP family regulates multiple pre- and post-transcriptional processes, including alternative splicing, transcript stability, RNA transport, and translational regulation [29],[31],[68],[69],[70],[71]. Importantly, direct regulation of hnRNP abundance by mTOR signaling has not been well-established in the literature except for one publication reporting mTOR binding to hnRNP F/H [67]. The coordinated restoration of multiple hnRNP family members that we observed in our studies, therefore, suggests a previously unrecognized link between mTOR signaling and RNA-binding protein networks in the cerebrovasculature. Our findings suggest that mTOR-mediated disruption of RNA-binding protein homeostasis may significantly contribute to cerebrovascular dysfunction in AD [20],[47].

Several hnRNPs have been linked to AD and related proteinopathies through changes in their abundance, mislocalization, or behavior in stress granules [63]. Importantly, prior studies showed that hnRNP E1 (HNRNP E1), binds to AU-rich elements within the 3′UTR of eNOS mRNA and regulates its stability and translation efficiency [39],[71],[72],[73]. Also, changes in hnRNP E1 levels have been shown to influence eNOS expression, nitric oxide production, and endothelial responses under inflammatory and oxidative stress [39],[71],[72],[73]. We found that restoration of hnRNP levels by mTOR inhibition restored hnRNP E1, suggesting a mechanistic link between mTOR-mediated regulation of RNA-binding proteins and the recovery of nitric oxide-dependent vascular function associated with mTOR inhibition, which we have previously demonstrated [20],[46],[47],[74]. Thus, mTOR-driven reduction of hnRNP D1 and hnRNP E1 in the microvasculature of mice modeling AD amyloidopathy (Fig. 1D, G) and tauopathy (Fig. 3C), and during disease progression in AD brains (Fig. 2A) may contribute to impaired brain microvascular endothelial dysfunction in AD [74],[75] and models of AD [5],[20],[46],[47],[74],[76],[77].

In our studies, hnRNP D1 and hnRNP E1 showed a biphasic, age-dependent pattern in our amyloidopathy model, hnRNP D1 and hnRNP E1 showed a biphasic, age-dependent pattern: an early, noncanonical elevation at 6 months that was not further increased by rapamycin for hnRNP D1 and was augmented by rapamycin for hnRNP E1, followed by a disease-associated deficit at 12 months that was partially restored toward WT levels by mTOR attenuation. This biphasic pattern is consistent with evidence that tau pathology disrupts RNA-binding protein homeostasis early in disease. Key RNA-binding proteins are increased in tauopathy mouse models and AD hippocampus, a response proposed to be initially compensatory [79]. Tau interactome studies further identify hnRNPs as early disease-associated tau interactors and describe stage-dependent dysregulation as tau pathology develops [78]. The later decline we observed is likewise consistent with reports that hnRNP D-like decreases during normal brain aging and declines further in an amyloid AD model [80].

These findings suggest that RNA-binding protein homeostasis in AD-like cerebrovasculature is dynamically regulated across disease stages: an early stress-associated or compensatory response may give way to a later loss of compensation as amyloidopathy advances. Rapamycin had different effects at these stages, indicating that the timing of mTOR-targeted intervention may be important. hnRNP D1 isoforms regulate mRNA stability and inflammatory-transcript decay through distinct mechanisms [81,82].

In human AD brain tissue, hnRNP E1 (r = −0.43, P = 0.039) and hnRNP L (r = −0.59, P = 0.005) were inversely associated with Braak stage, while hnRNP R and hnRNP M showed similar negative, non-statistically significant trends. hnRNP K and hnRNP A/B showed no association with Braak stage. These analyses were not adjusted for age, sex, PMI, or other covariates because the cohort was not powered for multivariable analyses. However together, the mouse and human data support selective disruption of an hnRNP RNA-binding protein network in AD, rather than a uniform loss of all hnRNP family members.

Importantly, our data suggest that pathogenic oligomeric tau in a model of tauopathy [56],[74],[83],[84],[85] is sufficient to downregulate hnRNP E1 in the microvasculature of this tauopathy model. These data can be interpreted in the context of the revised amyloid hypothesis [86], in which amyloid-β (Aβ) triggers pathological processes that lead to tau-mediated neurodegeneration [87]. Thus, vascular dysfunction arising from amyloid accumulation may be, at least in part, mediated by pathogenic tau through mTOR-driven downregulation of RNA-regulating protein networks. Chronic mTOR activation [88], linked to amyloid and tau pathology in humans, may thus suppress RNA regulatory pathways. Since hnRNPs control mRNA stability, alternative splicing, and endothelial stress-response transcripts, their depletion could weaken vascular resilience and increase susceptibility to ongoing amyloid- and tau-related damage. Restoring RNA-processing pathways and neurovascular function with rapamycin treatment further supports a role for mTOR-dependent RNA dysregulation in neurovascular impairment. Overall, our findings support a model in which vascular mTOR-mediated RNA regulatory dysfunction contributes to the pathogenesis and progression of AD and may be a therapeutically addressable factor in AD.

Limitations of the present studies include the lack of cell-type specificity afforded by the generation of microvascular fractions. While our studies suggest a role for HNRNP D1 alterations specifically in brain microvascular endothelial cells, future single-cell or spatial approaches will be necessary to identify the cell type(s) affected by mTOR-driven depletion of RNA-regulating proteins in AD. Additional mechanistic studies should determine whether restoring specific single hnRNPs can negate microvascular dysfunction in AD models of amyloidopathy and tauopathy. Despite these limitations, our studies provide a useful dataset reflecting the AD-like microvascular proteome in the presence and absence of mTOR attenuation and identify vascular RNA-binding protein homeostasis as a key regulator of neurovascular dysfunction across AD models. The present studies thus link vascular RNA regulatory disruption by mTOR activity to microvascular dysregulation in AD, broadening the mechanistic framework describing how vascular dysfunction contributes to AD progression. To our knowledge, our work provides the first evidence linking mTOR activity to the coordinated disruption of RNA-binding protein networks in the cerebrovasculature of AD and AD models and demonstrates that these changes can be reversed with interventions that inhibit mTOR.

## Acknowledgments

We acknowledge the following funding support: the National Institute on Aging (NIA) grant 1R01AG057964-01 (Galvan); the NIH/NIA Recovery Act Grand Opportunities grant RC2AG036613 (Richardson); the U.S. Department of Veterans Affairs, grant I01 BX002211-01A2 (Galvan); the Oklahoma Nathan Shock Center on Aging, NIA grant 2P30AG050911-11 (Galvan); the San Antonio Nathan Shock Center of Excellence in the Biology of Aging, NIA grant 2P30AG013319-21 (Galvan); the San Antonio Medical Foundation (Galvan); and the U.S. Department of Veterans Affairs, grant IK2 BX003798-01A1 (Hussong). Dr. Galvan acknowledges the generous support of the Robert L. Bailey and Daughter Lisa K. Bailey Alzheimer’s Fund, established in memory of Jo Nell Bailey, and the Protein Phenotypes of Aging Core at the University of Washington Nathan Shock Center 5P30AG013280-32 for mass spectrometry based proteomic studies and data analysis.

